# Substrate Binding and Inhibition of the Anion Exchanger 1 Transporter

**DOI:** 10.1101/2022.02.11.480130

**Authors:** Michael J. Capper, Shifan Yang, Alexander C. Stone, Sezen Vatansever, Gregory Zilberg, Yamuna Kalyani Mathiharan, Raul Habib, Keino Hutchinson, Avner Schlessinger, Mihaly Mezei, Roman Osman, Bin Zhang, Daniel Wacker

**Affiliations:** Department of Pharmacological Sciences, Icahn School of Medicine at Mount Sinai, New York, New York 10029; Department of Genetics and Genomic Sciences, Icahn School of Medicine at Mount Sinai, One Gustave L. Levy Place, New York, NY 10029, USA; Mount Sinai Center for Transformative Disease Modeling, Icahn School of Medicine at Mount Sinai, One Gustave L. Levy Place, New York, NY 10029, USA; Department of Neuroscience, Icahn School of Medicine at Mount Sinai, New York, New York 10029

## Abstract

Anion Exchanger 1 (AE1, SLC4A1) is the primary bicarbonate (HCO_3_^-^) transporter expressed in erythrocyte membranes where it mediates transport of CO_2_ between lungs and other tissues via import/export of bicarbonate. It is also a key regulator of erythrocyte structure and antigenic recognition. Previous biochemical studies, and a low-resolution crystal structure of the transmembrane domain have provided initial insight into AE1 structure and function. However, key questions remain regarding substrate binding and transport as well as the mechanism of inhibition. The orientation of the intracellular domain as well as the localization of lipid and sterol binding sites also remain enigmatic. We herein present seven novel high resolution cryo-EM structures of the full length human transporter in the apo, bicarbonate-bound, and several inhibitor-bound states combined with uptake- and computational studies. To our knowledge, these studies represent the first full length human, and substrate bound, SLC4 transporter structure. Our results reveal important molecular details about substrate binding and transport, as well as the diverse mechanisms of AE1 inhibition by both research chemicals and prescription drugs. We also provide novel insights into the full-length transporter architecture, identify the conformational space of the Diego blood antigen system and elucidate multiple lipid and sterol binding sites.

## Introduction

Anion exchanger 1 (AE1); or SLC4A1 is one of nine bicarbonate transporters in the SLC4 family of membrane proteins that help regulate cellular pH in virtually all tissues. AE1 is a key transporter in red blood cells (RBC) where it shuttles CO_2_ between lungs and other tissues via bicarbonate transport and contributes to the structural integrity of RBCs through interactions with the cytoskeleton. AE1’s extracellular surface binds antibodies of the Diego antigen system, a blood group of 21 antigens that can cause potentially fatal hemolytic disease of the newborn^1^: Antibodies produced by the mother can bind epitopes on AE1, thereby attacking RBCs of the fetus or the newborn^2^. Other mutations can disrupt AE1 structure and/or transport resulting in red blood cell deformities and kidney diseases such as renal acidosis^3,4^.

AE1 is modular, but the interplay between its cytoplasmic domain (cdAE1), which binds the cytoskeleton, and membrane domain (mdAE1), which mediates substrate transport, remains poorly understood. AE1 is thought to serve as a nucleus in the periphery of the RBC plasma membrane, forming complexes with enzymes and effectors involved in a variety of RBC biology^5,6^. mdAE1 is the functional site of electroneutral Cl^-^/HCO_3_^-^ exchange through an alternating access mechanism containing a single anion binding site, the precise nature of which is still unknown^3^. A diverse collection of chemical compounds, including clinical drugs, has been shown to inhibit AE1, but the distinct mechanisms by which these compounds inhibit AE1 have been limited to early NMR studies of substrate binding^4,7–10^.

Although several studies have provided insight into the roles of specific residues in the transport and inhibition of the transporter, key questions regarding inhibitory modes and global molecular mechanisms remain. Answers have been elusive due to the absence of structural insight into AE1, which has been limited to a single low-resolution crystal structure of mdAE1^11^. High-resolution cryo-EM has provided the opportunity to elucidate the bicarbonate binding site, as well as to characterize how diverse chemical inhibitors bind to and inhibit AE1. Herein, we report seven unique AE1 cryo-EM structures at resolutions ranging between 2.96-3.37 Å. Our studies allow for the unambiguous near-atomic characterization of the mdAE1 including the Diego antigens, bound lipids, sterols, substrates, and inhibitors, and provide insight into the interplay between the cytoplasmic and transmembrane domains of AE1.

## Results

### Structure Determination of full-length human AE1

Cryo-EM studies were carried out using the full length human AE1 purified from *Sf9* insect cells, and complexed with inhibitors, substrate, or in the “apo” state (see methods for details). Core areas of the structures reach local resolutions as high as 2.5 Å as calculated by local resolution estimation in cryoSPARC (Extended Data Figure 1, Extended Data Table 1). Our highest resolution structure of AE1 bound to the stilbene inhibitor DIDS (4,4’-diisothiocyanatostilbene-2,2’-disulfonic Acid) was obtained at 2.96 Å, and allowed us to identify several key features of the transporter not observed in the previous mdAE1 crystal structure^3,11^.

Overall, our structures conform with known SLC4 architecture; two AE1 protomers form a homodimeric complex (Fig 1). However, unlike previous SLC4 transporter structures^11–13^, our full-length AE1 reconstruction reveals the relative orientation of cdAE1 and mdAE1 (Fig 1, Extended Data Figure 1C). In our resulting model cdAE1 orients perpendicular to mdAE1. The lack of high-resolution information for the cdAE1 implies considerable flexibility between the domains as previously proposed^14^, but indicates that in the absence of binding partners, full-length AE1 protomers neither dimerize in a parallel nor a twisted orientation as has been debated^15^. Extensive 3D sub-classification was unable to produce sufficient density to build an atomic model of cdAE1, which further highlights the flexibility and dynamic nature of cdAE1 in the absence of binding partners. When refined without symmetry, a tilt was present in the cdAE1 that revealed non-covalent contacts between mdAE1 helices 5 and 6 and residues within the cdAE1 (Extended Data Fig 1C). Both, simulations^15^ and cross-linking experiments^16^ have proposed similar interactions, but our structure refutes both protomers forming a tight complex at the same time.

**Fig. 1.**
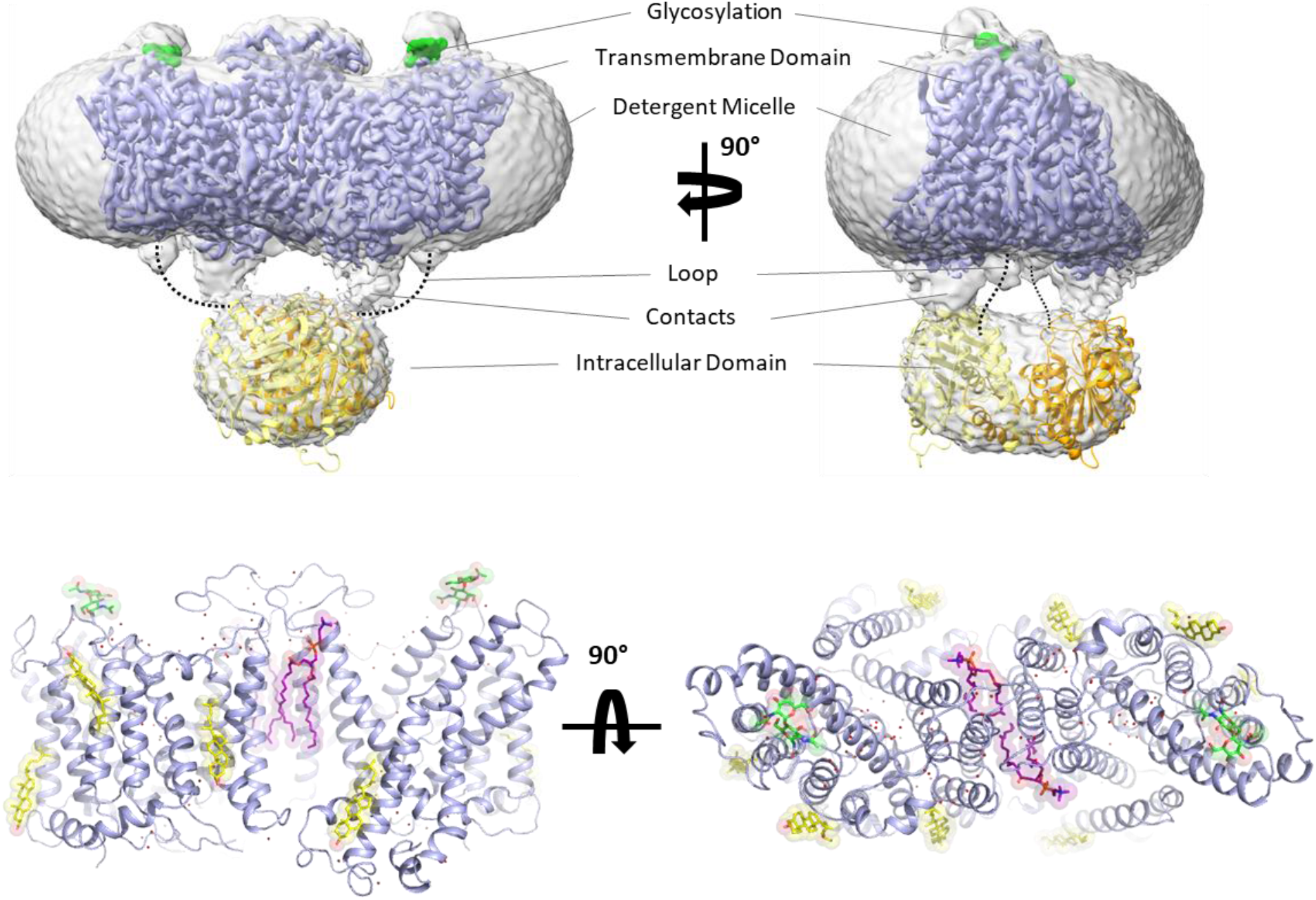
Cryo-EM structure of full length human AE1/SLC4A1. **Top**, Cryo-EM density of overall AE1 homodimer and detergent micelle (grey), overlaid with density of the membrane domain of AE1 (mdAE1, light blue). Glycosylation sites are highlighted in green, and the crystal structure of cytoplasmic domain (cdAE1) homodimer (yellow/orange) is loosely fit into density. Dotted lines highlight that the loop connecting cdAE1 and mdAE1 termini is in a different position from non-covalent contacts observed between the domains. **Bottom**, mdAE1 structure (light blue) including bound lipids (purple) and cholesterol (yellow), glycosylation sites (green), and water molecules (red spheres).

We further reveal the complete extracellular surface of the transporter (Fig 1) including all extracellular loop regions and glycosylation sites of both protomers. We were able to map the position and architecture of all Diego blood group antigens located on AE1 (Extended Data Figure 1D), revealing the detailed molecular architecture of the epitopes targeted in various hemolytic diseases^1,2,17^.

Lastly, we also observe density for multiple lipids and sterols in our structures. Some density remains unidentified, but we observe three cholesterol molecules per protomer in our highest resolution structure. One such cholesterol molecule is located at the interface of the gate and core domain (TM1 and TM7). Such binding could allosterically modulate the conformational changes required for transport and explain the inhibitory effect of cholesterol on AE1^18–20^. We also identify two phospholipids bound within the dimer interface between mdAE1 protomers, with the head groups interacting with the extracellular side of AE1 (Fig 1, Extended Data Figure 1).

### The Bicarbonate Binding Site

To better understand the substrate binding mechanism, we determined structures of “apo” and bicarbonate-bound AE1 (Fig 2). “Apo” refers to AE1 purified in the presence of 100 mM chloride, the structure of which does not show any density for chloride near the presumed anion binding site or elsewhere. In contrast, AE1 purified in the presence of 100 mM sodium bicarbonate in a chloride-free buffer (see methods), showed strong electron density attributed to bicarbonate (Fig 2, Extended Data Fig 2). Our near-atomic resolution structure is the first structure of any human SLC4 transporter to reveal the precise binding site of a substrate. In both “apo” and bicarbonate-bound structures, AE1 appears in the outward-facing state, with a channel-like cavity that is accessible from the extracellular site and leads to a positively charged cation selectivity filter/anion binding site near the ends of TM3 and TM10 (Fig 2)^3,21^. Reminiscent of how Uracil is bound in the SLC26 UraA transporter (Extended Data Fig 2C), density for bicarbonate is located near R730, which has previously been implicated in transport^3,11,21^. The negatively charged bicarbonate ion is bound in a small 23 Å^3^ pocket less than 3 Å from the R730’s side chain, indicating a strong ionic interaction. We observe weaker interactions between bicarbonate and backbone amide bonds in TM10, which was suggested as a positive dipole that can provide binding sites for anions^3^. Estimating the relative binding energy contributions of nearby residues in AMBER^22^ suggests that bicarbonate does not interact with residues at the N-terminal end of TM3, the other proposed dipole of the anion binding site. Moreover, while the R730 sidechain is the key anchor, backbone interactions with T727, T728, and V729 contribute substantially to bicarbonate binding (Extended Data Table 2). Using Simulated Annealing of Chemical Potential (SACP) simulations^23^ (Extended Data Fig 2D), we computationally estimate an apparent bicarbonate K_D_ of 1.6 mM for this site, which is similar to NMR studies that estimated a K_D_ of 5.4 mM^10^. To test the relevance of this affinity for AE1-mediated transport, we performed cellular bicarbonate uptake experiments and obtained a concentration of K=2.8 mM at which bicarbonate uptake reaches half-saturation (Fig 2G, see methods for details).

**Fig. 2.**
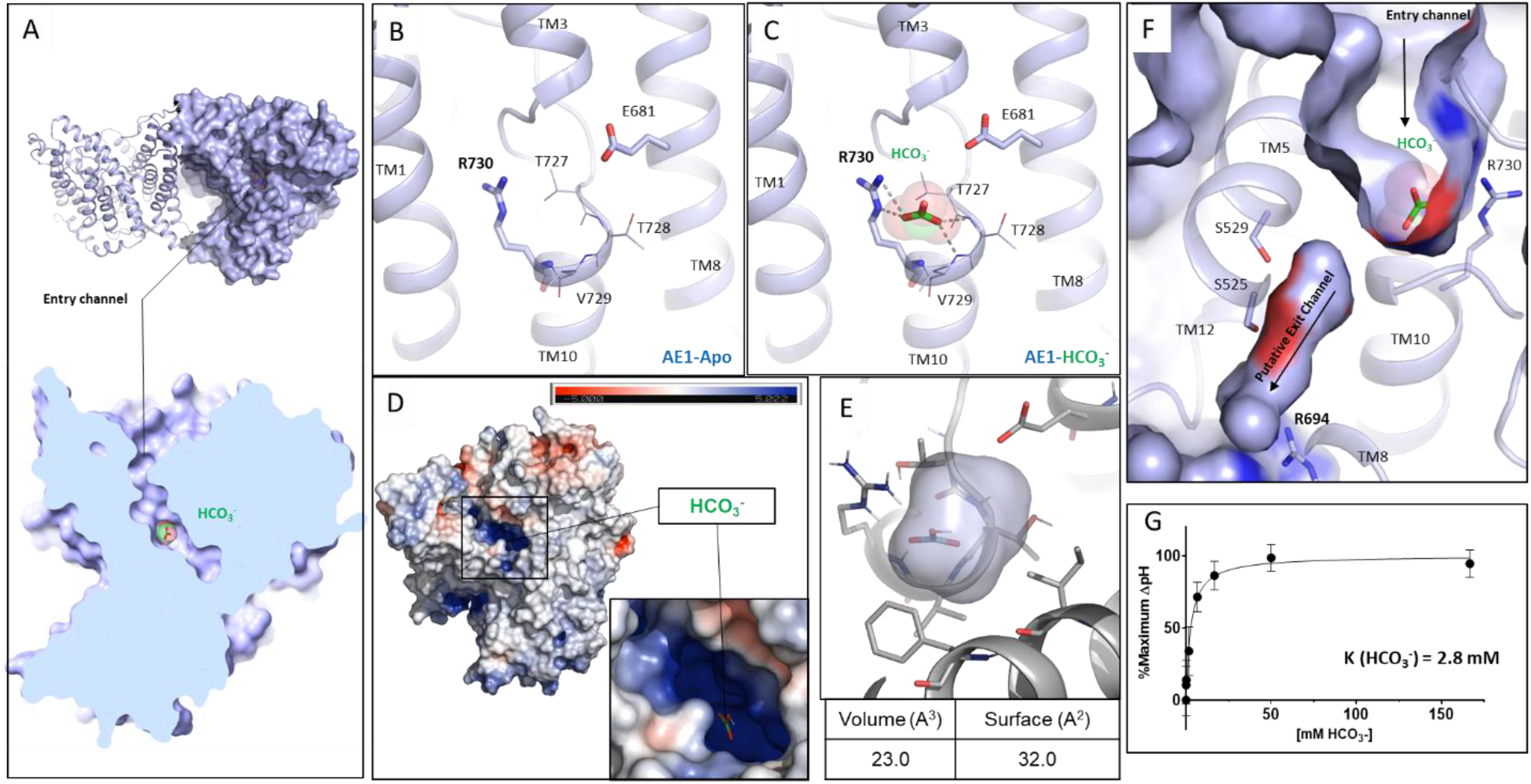
Structural insight into bicarbonate binding at AE1. **A**, Overall structure of mdAE1 homodimer (top), and cut site view of monomer showing bound bicarbonate (bottom, green). **B**, **C**, Close-up of anion binding site in apo and bicarbonate bound AE1 structures with highlighted key residues. **D**, mdAE1 charge distribution highlighting positively (blue) and negatively charged surfaces (red). **E**, Calculation of bicarbonate binding site volume and surface using POVME. **F**, Surface display shows putative anion exit channel leading from the bicarbonate binding site to the cytoplasmic site. **G**, Measurement of cellular bicarbonate uptake over one minute using different substrate concentrations. An uptake constant of K=2.8 mM was determined at which concentration half maximum uptake saturation was observed. Data are mean ± s.e.m. of three independent experiments (n=3) performed in triplicates, and were fit with a one-site saturation curve in GraphPad Prism.

Located just below the bicarbonate binding site within mdAE1, we observe a cavity formed by TM5, TM8, TM10, and TM12 between the core and gate domains of AE1 (Fig 2). This cavity is lined with two serines and constrained by R694 at the cytoplasmic side. These properties and the proximity to bicarbonate suggest that this cavity could expand to become part of a putative substrate exit tunnel in an inward-facing state. In fact, our SACP simulations identify a second bicarbonate binding site in which the anion binds to R694 and S525 (Extended Data Fig 2D), providing further evidence for this proposed exit path.

### Molecular mechanisms of transport inhibition

To investigate the structural basis of how different compounds inhibit SLC4-mediated substrate transport, we next determined several structures of AE1 bound to different inhibitors including two clinical drugs.

#### Competition for substrate binding

The stilbene compound H_2_DIDS (4,4’-Diisothiocyanatodihydrostilbene-2,2’-Disulfonic Acid) was used in the previously published mdAE1 structure and appears to be covalently linked to both K539 in TM5 and K851 in TM13. According to previous studies, both lysines are covalently bound at 37°C and pH 9.5, while lower pH prevented linkage of K851^24^. Due to some poorly defined electron density in the lower resolution mdAE1-H_2_DIDS crystal structure^11^, there indeed remains some ambiguity regarding the bond with K851. We thus investigated transporter binding by stilbene inhibitors and determined sub 3 Å structures of AE1-DIDS and AE1-H_2_DIDS formed under lesser alkaline conditions (pH 9, 22°C). Our structures show covalent binding to K539 only, while K851 appears to form ionic interactions with the stilbene’s sulfonic acid group and E535 (Fig 3, Extended Data Fig 3). We thus reason that harsher conditions than we used are required to weaken these interactions and facilitate covalent binding to K851.

**Fig. 3.**
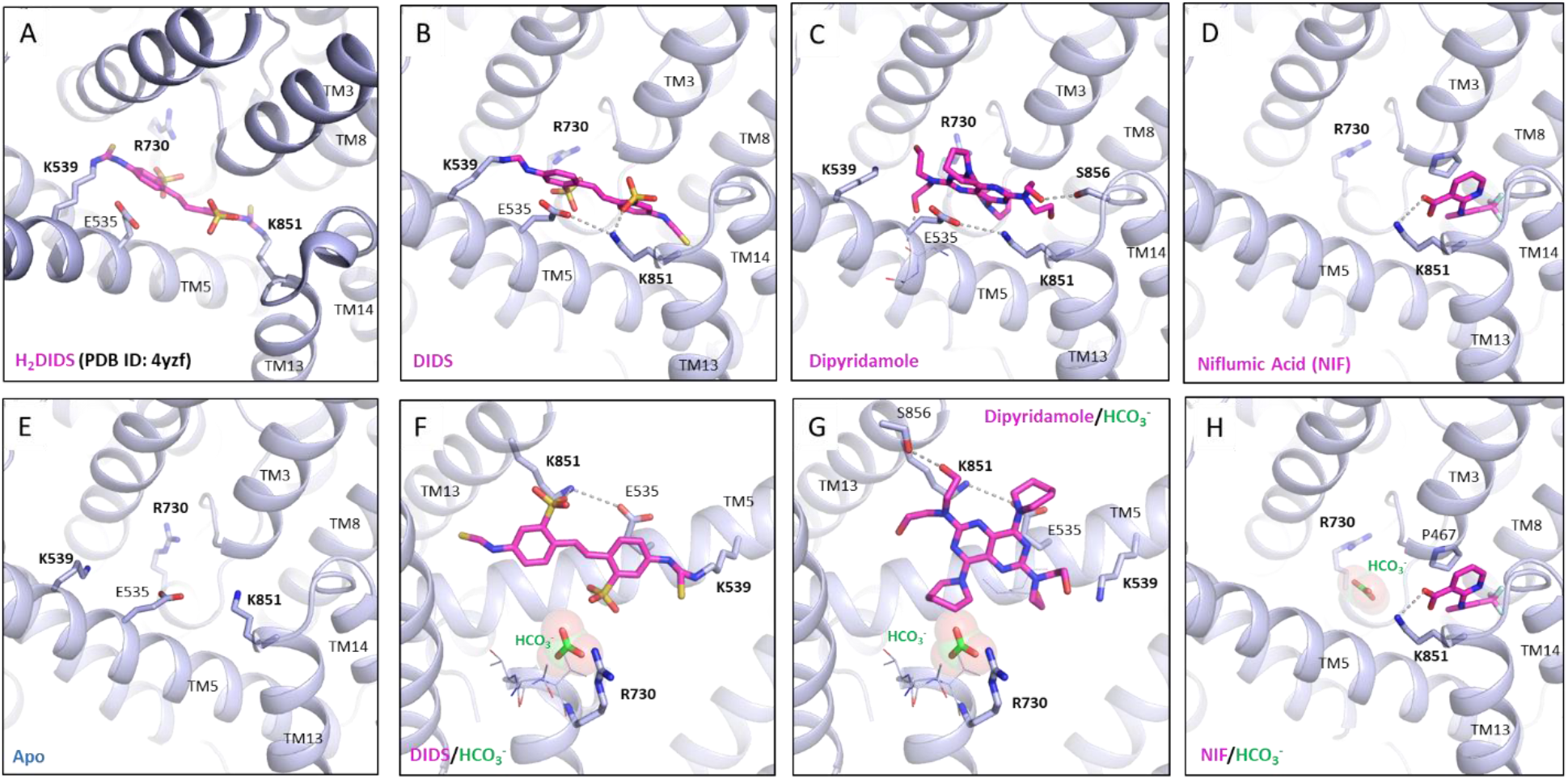
Structure of AE1 bound to chemically and pharmacologically diverse inhibitors. **A**, Previous mdAE1-H_2_DIDS crystal structure (PDB ID: 4YZF) showing likely covalent binding of H_2_DIDS (magenta) to K539 and K851. Cryo-EM structures of AE1 bound to DIDS (**B**), Dipyridamole (**C**), NIF (**D**), or Apo (**E**) reveal the binding location and pose of inhibitors (magenta). Overlay with bicarbonate (green) bound AE1 structure shows that DIDS (**E**) and Dipyridamole (**F**) restrict access to anion binding site, while NIF (**G**) binds in a different location and leaves access to anion binding site unobstructed. Ionic interactions and hydrogen bonds are shown as dotted lines.

When compared with bicarbonate-bound AE1, both DIDS and H_2_DIDS are located in the access channel leading from the extracellular space to the buried anion binding site. We further observe that one of the stilbene’s sulfonic acid groups is located less than 4 Å from the bicarbonate ion (Fig 3F). These findings indicate that DIDS/H_2_DIDS not only block access to the anion binding site but likely also affect bicarbonate binding through charge repulsion as there remains sufficient space to bind ions. Our findings provide a structural explanation for NMR studies that showed that DIDS reduces substrate affinity. It should be noted that several previous studies investigated the substrate Cl^-^ not bicarbonate, but competition of both anions for the same site indicates a common or similar binding site^10^.

We next determined a 3.07 Å structure of AE1 treated with diethyl pyrocarbonate (DEPC) (Extended Data Figure 3), which has been reported to inhibit transport and stilbene binding by stabilizing an inward-facing conformation via covalent modification of H834^25,26^. Surprisingly, our AE1-DEPC structure shows an outward-facing state, with no electron density accounting for a modified H834 side chain. Instead, we observe that DEPC covalently modifies K539 and K851 (Extended Data Fig 3F-G), indicating that modification of K851 not H834 leads to the increased mass of an AE1 fragment in previous work^25^. We, therefore, suggest that DEPC-mediated modification of K851 sterically precludes H_2_DIDS binding rather than stabilizes an inward-facing state^26^.

#### Inhibition through substrate channel blocking

While DIDS reduces anion affinity^7^, other inhibitors have been described to only block access to the transport site^8^. One such inhibitor is the FDA-approved antiplatelet medicine dipyridamole, which has been shown to block AE1 substrate channels whilst not competing for anion binding^8^. To elucidate the different mechanisms by which stilbenes and dipyridamole inhibit transport, we determined a 3.19 Å structure of AE1-dipyridamole (Fig 3, Extended Data Fig 3D). This structure reveals that the drug occupies a similar site as DIDS and H_2_DIDS in the same outward-facing transporter conformation. Specifically, dipyridamole stretches between the core and gate domain, where it forms hydrogen bonds with a backbone carbonyl in TM5 and S856 of TM3. One of the piperidine rings appears to stack in the dipole region between TM3 and TM10 towards E681 and closer to TM3. However, dipyridamole binding lacks the charge repulsion provided by the sulfonic acid group of the stilbene compounds, which likely explains why the compound does not compete for anion binding^8^.

#### Inhibition of translocation

Niflumic Acid (NIF), an analgesic and anti-inflammatory drug used in the treatment of rheumatoid arthritis, is a chloride channel inhibitor that has previously been shown to inhibit AE1 substrate transport through a mechanism distinct from DIDS, H_2_DIDS, or dipyridamole^9^. Specifically, studies have shown that NIF does not affect substrate affinity or access to the binding site, but instead inhibits transport by preventing transition between outward- and inward-facing states^9^. To investigate the molecular basis for this distinct pharmacology, we determined a 3.18 Å cryo-EM structure of AE1-NIF. Consistent with a different mechanism of action, we observe NIF bound to a different site than H_2_DIDS, DIDS, and dipyridamole (Fig 3, Extended Data Figure 4A-E). NIF appears to be accommodated in a 138 Å^3-^ sized pocket between the core and gate domains formed by TM3, TM8, TM13, and TM14, which overlaps only partially with dipyridamole’s binding pose and the non-attached isothiocyanate groups of DIDS/H_2_DIDS (Fig 3). We also performed molecular docking studies, which further validated NIF’s unexpected binding pose and location (Extended Data Fig 4C). NIF appears to be anchored by a salt bridge between its carboxylate group and K851. In addition, the compound is wedged tightly between P467 in TM3 and L859 in TM14 causing structural rearrangements to accommodate the compound. We note subtle outward movements of the solvate exposed tips of TM13 and TM14, as well as changes in e.g., F524, L859, and K851 (Extended Data Fig 4D). Despite inhibiting transport akin to DIDS/H_2_DIDS (Extended Data Fig 4F), NIF however does not seem to obstruct access to the bicarbonate binding pocket (Fig 3H). Our structure thus suggests that NIF binding between the AE1 gate and core domains prevent translocation-related changes, while not interfering with substrate binding^9^.

## Discussion

We herein report seven novel high-resolution cryo-EM structures of the full-length human AE1 transporter bound to substrate and multiple different drugs and therapeutics. We provide key insights into the transporter architecture, fully elucidate the Diego blood group antigens associated with severe hemolytic diseases, and identify lipid and sterol binding sites. Compared to previous work, our studies reveal a fully ordered extracellular surface, including all Diego blood group antigens and glycosylation sites. This observation is surprising as AE1’s extracellular surface does not contain secondary structure elements compared to e.g., NDCBE (SLC4A8)^13^. We also reveal several bound lipids and sterols, whose effects on AE1 structure and function have been heavily speculated on^19,27,28^. Contrasting previous studies that propose cholesterol at the dimer interface, we only observe cholesterol bound on membrane-facing surfaces^19^. But we do observe lipids bound to the dimer interface which has been proposed to stabilize and regulate the structure-function of the transporter^19^.

At global resolutions of up to 2.96 Å, we also illuminate in near-atomic detail how the substrate bicarbonate binds to AE1, and structurally characterize how chemically and pharmacologically distinct inhibitors differentially affect both substrate binding and transport (Fig 4). Based on structures, uptake assays, and computational studies we propose that R730 forms the center of the anion binding site and holds the anion in place with low millimolar affinity before conformational transition and substrate translocation. Our observed bicarbonate binding site is similar to that identified in computational modelling^29^, but it is distinct from that of a related sodium-dependent bicarbonate transporter NDCBE (SLC4A8)^13^. However, we have limited structural comparison due to this structure’s problematic ion placement with several observed clashes and suboptimal bicarbonate residue environment - likely a result of the lower resolution and ambiguous electron density (Extended Data Fig 2E-F).

Analysis of our structures in the context of other transporter structures provides intriguing insights into AE1’s transport mechanisms, which have remained largely elusive. When compared to a previous SLC4 borate transporter structure from *A. thaliana* (AtBor1)^30^ our findings strongly indicate that AE1 transports bicarbonate via an elevator mechanism (Fig 4), as hypothesized in previous studies^31^. The well-ordered EL3 and lipids facilitating dimerization^19^ argue for a stationary gate domain that is consistent with an elevator mechanism, but not e.g. a rocker-switch model^32^, and has been described as a common feature of oligomeric elevator transporters^33^. Superposition of AE1 with the AtBor1 structure shows that the gate domains align well, while the AtBor1 core domain appears shifted downward (Fig 4B). TM3 and TM10, which form the AE1 bicarbonate binding site, move downwards by about 5-7 Å, and TM10 bends away from the gate domain, thus likely releasing bound substrate towards the intracellular site. This is further supported by a cytoplasmic exit channel in AtBor1 that connects to the AE1 bicarbonate binding site even before translocation of AE1’s core domain. A cavity observed in our AE1 structures (Fig 2F) overlaps well with this channel, and likely expands into a substrate exit channel during substrate translocation. In fact, our studies suggest that R694 located at the cytoplasmic exit could form a second bicarbonate binding site. Further evidence for this transport model comes from a cholesterol bound between the core and gate domain at the interface of TM1 and TM7 (Fig 1), which may prevent translocation-related conformational changes and explain the inhibitory effects of cholesterol^18,20^. Similarly, although NIF does not appear to inhibit bicarbonate binding, its location between the core and gate domain likely prevents relative domain movements required for ion transport. These findings together with the proposed shared substrate binding site^10^ suggest a general exchange mechanism in which bicarbonate or chloride is transported one way, before the counterion is transported the opposite way and AE1 is returned to its resting state.

**Fig. 4.**
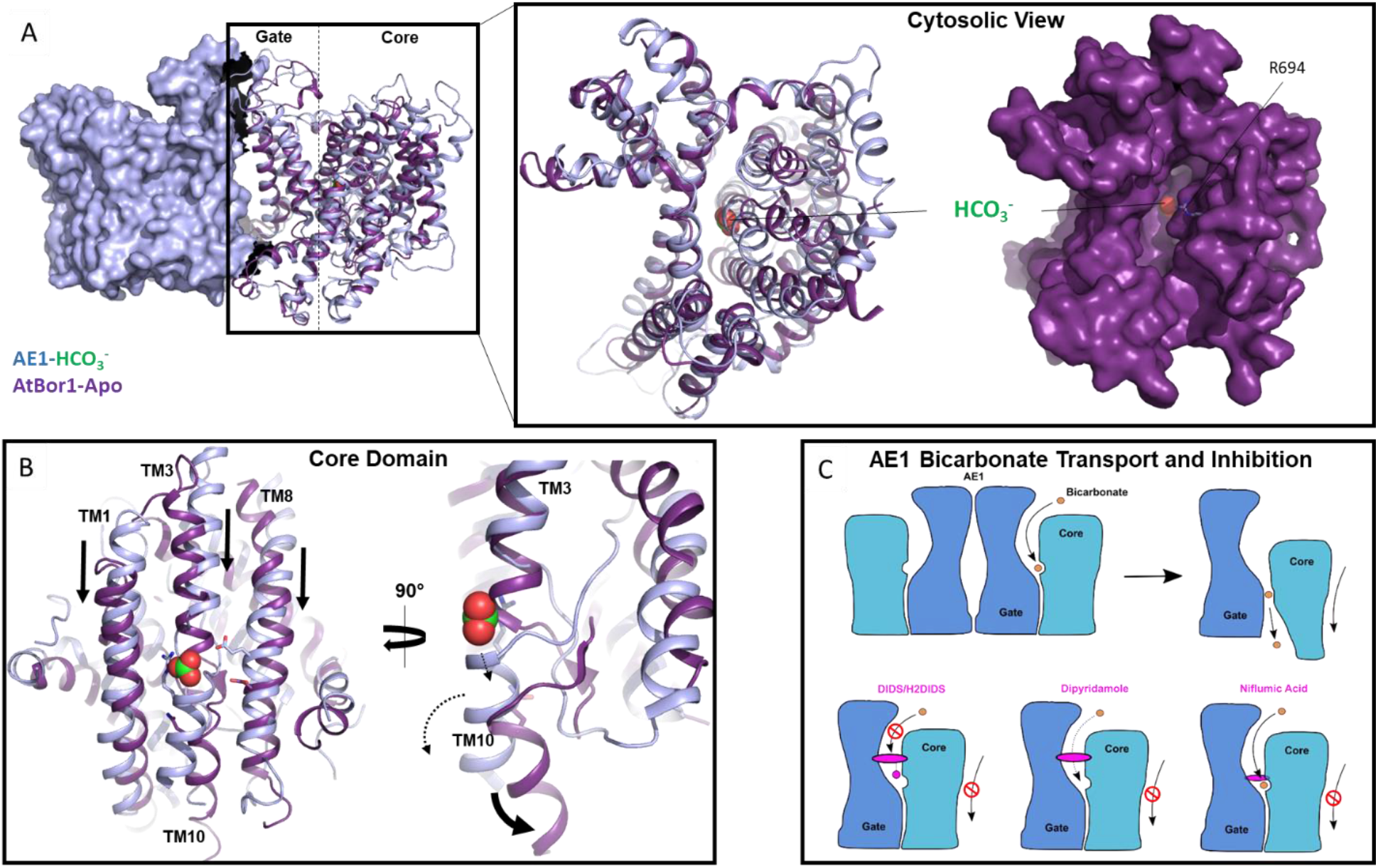
Model of AE1-mediated bicarbonate transport and diverse mechanisms of AE1 transport inhibition by pharmacologically different drugs. **A**, Overlay of membrane domains of human AE1 (light blue) and AtBor1 from *A. thaliana* (purple) showing similar gate conformation, while the core domains appear in different states. Cytoplasmic view of the overlay reveals a channel in AtBor1 that overlaps with the putative anion exit channel in AE1 ending near R694, and connecting the AE1 bicarbonate binding site and the cytosol. **B,** Overlay of AE1 and AtBor1 anion binding sites suggests an elevator mechanism where the core domain moves towards the cytoplasmic site (black arrows) and TM10 kinks away from the gate domain (black arrow) to release substrates towards the cytoplasm (dashed arrows). **C,** Schematic illustrating AE1-mediated bicarbonate transport, as well as pharmacological differences between the inhibitors DIDS/H_2_DIDS, Dipyridamole, and Niflumic Acid. AE1 gate and core domains are shown in blue and teal, respectively, bicarbonate is shown in orange, and inhibitors are colored magenta.

The herein presented work thus not only reveals fundamental mechanisms of AE1-mediated substrate binding and transport, but also illuminates the different pharmacological mechanisms by which distinct research compounds and clinically used drugs such as Niflumic acid and dipyridamole inhibit transport. In fact, dipyridamole increases red blood cell deformability^34^ thus facilitating healthy red blood cell circulation, an effect likely mediated by AE1’s role in shaping RBC structure^35^. Given AE1’s physiological importance in RBC-mediated CO_2_ transport and acid secretion in the kidney, our structural insights should greatly facilitate the design of novel pharmacological tools to study distal renal tubular acidosis, hemolytic anemias, and other AE1-associated pathologies. Moreover, due to the similar molecular mechanisms of the SLC4, SLC26, and SLC23 anion transporter families, our findings will also inform mechanistic studies of other important transporters in human health and disease.

## Methods

### Construct and Expression

Structural studies reported herein were performed with the full-length human AE1 transporter (UniprotKB-P02730), which was cloned into a modified pFastBac vector to introduce a C-terminal 3C protease cleavage site followed by a 10xHis tag. Bacmid DNA was generated in DH10Bac cells (Invitrogen) and protein was expressed in *Sf9* cells (Expression Systems) using the Bac-to-Bac Baculovirus expression system (Invitrogen). ~2.5 μg recombinant bacmid DNA and 3 μl FuGENE HD Transfection reagent (Promega) in 100 μl Sf900 II media (Invitrogen) were added to 500,000 *Sf9* cells plated in 2 ml of SF900 II media in wells of a 12-well plate. After 5 days at 27 °C the supernatant was harvested as P0 viral stock, and high-titer recombinant P1 baculovirus (>10^9^ viral particles per ml) was obtained by adding 200 μl P0 to 40 ml of 3 ×10^6^ cells/ml and incubating cells for 3 days while shaking at 27 °C. Titers were determined by flow cytometric analysis staining P1 infected cells with gp64-PE antibody (Expression Systems). Expression of AE1 for structural studies was carried out by infection of *Sf9* cells at a cell density of 2-3 ×10^6^ cells/ml with P1 virus at MOI (multiplicity of infection) of 5. After 48 hrs of shaking at 27 °C, cells were harvested by centrifugation at 48 h post-infection and stored at −80 °C until use.

### Protein Purification and grid preparation

Typically, we purified protein from ~ 3 L of expression culture to prepare grids for cryo-EM experiments. Insect cell membranes were disrupted by thawing frozen cell pellets in a hypotonic buffer containing 10 mM HEPES pH 7.5, 10 mM MgCl2, 20 mM KCl and home-made protease inhibitor cocktail (500 μM AEBSF, 1 μM E-64, 1 μM Leupeptin, 150 nM Aprotinin) (Gold Biotechnology). Total cellular membranes were harvested by ultracentrifugation, and extensively washed by repeated (2-4 times) homogenization and centrifugation in a high osmotic buffer containing 1 M NaCl, 10 mM HEPES pH 7.5, 10 mM MgCl2, 20 mM KCl and home-made protease inhibitor cocktail. Purified membranes were directly flash-frozen in liquid nitrogen and stored at −80 °C until further use.

Purified membranes were resuspended in buffer containing 10 mM HEPES pH 7.5, 10 mM MgCl2, 20 mM KCl, 150 mM NaCl, home-made protease inhibitor cocktail, and 25 μM DIDS or H_2_DIDS, or 100 μM Dipyridamole or Niflumic Acid for the different AE1-inhibitor complexes. Complexation was initiated by agitation for 1 hr at room temperature, a step that was skipped for the AE1 “apo”, AE1-bicarbonate and AE1-DEPC samples. Prior to solubilization, membranes were equilibrated at 4 °C and incubated for 30 min in the presence of 2 mg/ml iodoacetamide (Sigma). Membranes were then solubilized in 10 mM HEPES, pH 7.5, 150 mM NaCl, 1% (w/v) n-dodecyl-β-D-maltopyranoside (DDM, Anatrace), 0.2% (w/v) cholesteryl hemisuccinate (CHS, Anatrace), inhibitor, and home-made protease inhibitor cocktail for 2 h at 4 °C. Unsolubilized material was removed by centrifugation at 200,000 × g for 30 min, and buffered imidazole was added to the supernatant for a final concentration of 20 mM. Proteins were bound to TALON IMAC resin (Clontech) overnight at 4 °C. Purification of the Dipyridamole and NIF bound complex was carried out in the presence of 50 μM inhibitor and the bicarbonate bound complex was purified in the presence of 100 mM sodium bicarbonate. The resin was then washed with 10 column volumes (cv) of Wash Buffer I (25 mM HEPES, pH 7.5, 500 mM NaCl, 0.1% (w/v) DDM, 0.02% (w/v) CHS, 20 mM imidazole, 10% (v/v) glycerol). The detergent was then exchanged for LMNG by successively incubating the resin with the following buffers for 1 hour each: Wash Buffer II (25 mM HEPES, pH 7.5, 500 mM NaCl, 0.05% (w/v) DDM, 0.05% (w/v) LMNG, 0.02% (w/v) CHS), Wash Buffer III (25 mM HEPES, pH 7.5, 500 mM NaCl, 0.025% (w/v) DDM, 0.075% (w/v) LMNG, 0.02% (w/v) CHS), Wash Buffer IV (25 mM HEPES, pH 7.5, 500 mM NaCl, 0.05% (w/v) LMNG, 0.02% (w/v) CHS), Wash Buffer V (25 mM HEPES, pH 7.5, 500 mM NaCl, 0.025% (w/v) LMNG, 0.02% (w/v) CHS). After the final incubation step, the proteins were eluted with 25 mM HEPES, pH 7.5, 500 mM NaCl, 0.025% (w/v) LMNG, 0.02% (w/v) CHS and 250 mM imidazole. Protein purity and monodispersity were tested by SDS-PAGE and analytical size-exclusion chromatography (aSEC). Typically, the protein purity exceeded 95%, and the aSEC profile showed a single peak, indicative of transporter monodispersity. For the AE1-DEPC sample, we then added 5 mM DEPC and incubated the sample overnight at 4 °C. All complexes were finally purified over a S200 size exclusion chromatography column equilibrated in 20 mM HEPES, pH 7.5, 150 mM NaCl, 0.0011% (w/v) LMNG, 0.00011% (w/v) CHS, 0.00025% GDN. For AE1 bound to NIF and Dipyridamole, 50 μM of the respective compound was added to the buffer. The bicarbonate complex was purified in 20 mM HEPES, pH 7.5, 100 mM NaCHO_3_ 0.001% (w/v) LMNG, 0.0001% (w/v) CHS, 0.0001% GDN. Peak fractions were then pooled, concentrated to ~3-7 mg/ml, and immediately used to prepare grids for cryo-EM data collection.

### Grid Preparation, Cryo-EM Data collection and Processing

To prepare cryo-EM grids for imaging, 3 μl of purified AE1-Apo at ~6.3 mg/ml, AE1-bicarbonate at 5 mg/ml, AE1-DIDS at ~5 mg/ml, AE1-H_2_DIDS at ~4.1 mg/ml, AE1-DEPC at 5 mg/ml, AE1-Dipyridamole at 4.8 mg/ml, or AE1-Niflumic acid at 5 mg/ml were applied to glow-discharged holey carbon EM grids (Quantifoil 300 copper mesh, R1.2/1.3) in an EM-GP2 plunge freezer (Leica). EM-GP2 chamber was set to 95% humidity at 12°C. Sample-coated grids were blotted for 3 to 3.3 seconds before plunge-freezing into liquid ethane and stored in liquid nitrogen for data collection.

All automatic data collection was performed on a FEI Titan Krios equipped with a Gatan K3 direct electron detector run and operated by the Simons Electron Microscopy Center in the New York Structural Biology Center (New York, New York) or the Laboratory of BioMolecular Structure at Brookhaven National Laboratory. The microscope was operated at 300 kV accelerating voltage, at a nominal magnification of 64,000-81,000 corresponding to a pixel size of 1.08 Å. For each dataset, at least 3,500 movies were obtained at a dose rate of 25-30 electrons per Å^2^ per second with a defocus ranging from −0.5 to −1.8 μm. The total exposure time was 2 s and intermediate frames were recorded in 0.05 s intervals, resulting in an accumulated dose of 50-60 electrons per Å^2^ and a total of 40 frames per micrograph.

Movies were motion-corrected using MotionCor2^36^ and imported to cryoSPARC for further processing^37^. For CTF estimation we used patchCTF in cryoSPARC. An initial model was produced using a subset of micrographs and manual picking. Subsequent models were produced from particles found using templates. Datasets were curated by the removal of micrographs deemed irredeemable by poor CTF estimation. Particles were subject to 2D classification which quickly identified both the mdAE1 and cdAE1. A good initial model of mdAE1 was generated using ab-initio model building in cryoSPARC as were several bad models from rejected particles as a sink in future hetero-refinement. Multiple rounds of hetero-refinement were carried out to improve resolution followed by NU-refinement^37,38^. Stuctures were then further refined in ServalCat^39^, and final maps were generated in PHENIX^40^ before import into PyMOL^41^ for generating figures shown in the manuscript.

### Bicarbonate Transport Assay

A polyclonal HEK cell line that stably expresses AE1 upon tetracycline induction was generated using the TRex system (Invitrogen). Cells were plated in a 96-well plate and incubated overnight with or without 2 μg/ml tetracycline at 37° C. The next day, induced cells were again incubated with tetracycline for 3-4 hours. Cellular bicarbonate uptake was then determined via cellular changes in pH as previously described for other SLC4 transporters^12^. Cells were loaded with 5 μM of the pH-sensitive fluorescent dye BCECF-AM (2’,7’-Bis-(2-Carboxyethyl)-5-(and-6)-Carboxyfluorescein, Acetoxymethyl Ester) for 30 minutes. Following another short incubation in Hank’s balanced salt solution (HBSS) buffered with 50 mM HEPES pH 7.5, intracellular fluorescence ratio (excitation 495±20 nm and 435±20 nm; emission 540±30 nm) was measured using a multimode plate reader (Victor NIVO, Perkin Elmer). To initiate uptake, cells were then diluted 1:3 in Cl-free buffer (50 mM HEPES pH 7.5 adjusted with NaOH, 115 mM Na gluconate, 2.5 mM K_2_HPO_4_, 7 mM Ca gluconate, 1 mM Mg gluconate, 5 mM glucose, 30 μM amiloride), supplemented with different NaHCO_3_ concentrations. Fluorescence was then measured after 1 min. A calibration experiment using 10 μM nigericin in modified HBSS (1.26 mM CaCl_2_, 0.493 mM MgCl_2_, 0.407 mM MgSO_4_, 140 mM KCl, 0.441 mM KH_2_PO_4_, 4.17 mM NaHCO_3_, 0.338 mM Na_2_HPO_4_, 10 mM HEPES) at a range of pH values between 7 and 8 was then performed to convert fluorescence to pH values^42^.

Due to challenges associated with determining V_max_ (AE1 is reportedly the fastest human transporter with up to 10^5^ ions per second^3^) and cellular changes associated with changing pH, we opted not to determine a K_M_. Instead, we set out to determine an uptake constant via a saturatable uptake assay, which was performed in induced and uninduced cells using 0-167 mM NaHCO_3_. Specific uptake was then determined by plotting pH differences between induced and uninduced cells in response to different NaHCO_3_ concentrations. This was done in GraphPad Prism using a one-site saturation curve and allowed calculation of an uptake constant K=2.8 mM at which NaHCO_3_ concentration pH differences between induced and uninduced cells were at half maximum.

To measure AE1 inhibition, this experiment was performed in induced and uninduced cells using 16.7 mM NaHCO_3_. For DIDS and H_2_DIDS, cells were preincubated with 20 μM inhibitor in HEPES-buffered HBSS for 1 hr, after which DIDS and H_2_DIDS were omitted from the experiment. For NIF, 50 μM were added throughout the experiment after dye loading. Dipyridamole has spectral overlap with BCECF-AM and could therefore not be included in our measurements. All experiments were performed in triplicates, and data was averaged from three independent experiments and is shown as mean ± s.e.m.

### MD simulations

Even though AE1 exist in a dimeric form, it appears that the functional properties of each monomer are independent of the other. Thus only one monomer was selected for the construction. The system was built with CHARMM-GUI^43^ adding two cholesterol molecules and one cholesteryl succinate in the positions identified in the cryo-EM structure. The membrane was constructed from 200 POPC in both layers divided to account for the different surface area of the protein in the upper and lower leaves. The concentration of neutralizing K^+^ and Cl^-^ counterions in the rectangular box was set to ~0.15 mM. The initial apo-AE1 structure was translated using the charmm2lipid routine in AMBER^22^, and the simulations were conducted in AMBER 20. The system was minimized and equilibrated with the restraints designed in CHARMM-GUI. At the end of the equilibration, the MD simulations were executed at NPT conditions for 1000 ns. The final trajectory included 10,000 structures. A similar design and simulations were performed on the bicarbonate-occupied structure. The analysis of the trajectories were performed with cpptraj in AMBER and the simulaid facility^44^. The RMSD of the protein stabilizes after 100 ns and remains nearly constant for the rest of the simulation.

### Molecular Docking

To characterize the binding mode of NIF at AE1 using molecular docking calculations, we first removed the ligand from the cryo-EM structure of the AE1-NIF complex. AE1 was then prepared with the Maestro Protein Preparation Wizard under default parameters^45^. The binding site was defined by generating a grid with the Receptor Grid Generation Panel. The binding site outlining box was defined around the reference NIF ligand in the AE1 template structure. The NIF compound structure was obtained from PubChem (PubChem CID: 4488), and it was prepared for docking using LigPrep with the default parameters, where the possible states were generated at target pH 7±2. Docking was performed using Glide from the Schrödinger suite (Schrödinger, 2021). Finally, we used molecular mechanics generalized with born surface area solvation (MM-GBSA) with Prime in the Schrödinger suite to estimate the relative binding affinity between NIF and AE1^46^, where a more negative value of ΔG binding indicates higher binding affinity.

### Calculation of Bicarbonate Binding Energies

To compute the binding energy of bicarbonate we took the approach of Simulated Annealing of Chemical Potential (SACP)^23^. Briefly, the system is placed in a periodic box, which is divided into an inner box whose dimensions are 10 Å beyond the boundaries of the molecule and an outer (“bulk”) box of additional 5Å thickness. Using a Grand Canonical Ensemble/Monte Carlo (GC/MC) approach the entire system is equilibrated with inserting/deleting bicarbonate to reach a density of 0.15 g/L. The B parameter^47^, which reflects the excess chemical potential (B = μ_εX_ + ln<N>) in the “bulk” box is then decreased progressively. The change in the B parameter increases the probability of deletion of bicarbonate until the system equilibrates. The value of the B parameter at the point where the last bicarbonate is deleted equals the most negative energy of bicarbonate to the protein. An analysis of the B value at which bicarbonate is most proximal to a specific site (e.g., R730) yields the affinity of bicarbonate to this site. To enhance the statistical significance of the computed values, the MD trajectory was divided into 10 clusters and the center of the cluster was extracted to perform the SACP on each of them. The final result is the population weighted average of all the clusters for a specific location of the bicarbonate.

### Pocket Volume Analysis

POVME3^48^ (Pocket Volume Measurer 3) was used to calculate binding site volumes. We used default parameters for ligand-defined inclusion region, using the recently resolved structures as input PDBs. Pocket volume was visualized using PyMOL^41^.

## Acknowledgements

This work was supported by NIH grant GM133504, a Sloan Research Fellowship in Neuroscience, an Edward Mallinckrodt, Jr. Foundation Grant, a McKnight Foundation Scholars Award (all to D.W.), NIH T32 Training Grant GM062754 (G.Z.), R01 GM108911 (A.Sch. and K.H.), NIH Grant U01AG046170 (B.Z.), NIH Grant RF1AG057440 (B.Z.), NIH Grant R01AG068030 (B.Z.). NIH Grants R01DK073681, R01DK067555, R01DK061659 (R.O). Some of this work was performed at the National Center for cryo-EM Access and Training (NCCAT) and the Simons Electron Microscopy Center located at the New York Structural Biology Center, supported by the NIH Common Fund Transformative High Resolution Cryo-Electron Microscopy program (U24 GM129539,) and by grants from the Simons Foundation (SF349247) and NY State Assembly. We further acknowledge cryo-EM resources at the National Resource for Automated Molecular Microscopy located at the New York Structural Biology Center, supported by grants from the Simons Foundation (SF349247), NYSTAR, and the NIH National Institute of General Medical Sciences (GM103310) with additional support from Agouron Institute (F00316) and NIH (OD019994). For additional data collection we are also grateful to staff at the Laboratory for BioMolecular Structure (LBMS), which is supported by the DOE Office of Biological and Environmental Research (KP160711). This work was supported in part through the computational resources and staff expertise provided by Scientific Computing at the Icahn School of Medicine at Mount Sinai. We also like to thank J. F. Fay for help with initial data processing.

## Author Contributions

M.J.C. designed experiments, expressed and purified protein for grid freezing, collected data, refined structures, and helped write the manuscript. S.Y. and A.S. purified protein, prepared samples for grid freezing, and performed functional assays. S.V. performed computational studies and helped analyze the structures. G.Z. prepared grids for structure determination and assisted with data collection. Y.K.M. helped with data processing and structure refinement. R.H. helped establish protein expression and purification. S.V, K.H. performed docking studies and volume calculations supervised by A.Sch. R.O. performed molecular simulations and SACP analysis of substrate binding with help from M.M. B.Z. contributed to the study design and supervised computational studies. D.W. designed experiments, analysed the data, supervised the overall project and management, and wrote the manuscript.

## Data Availability

Density maps and structure coordinates have been deposited in the Electron Microscopy Data Bank (EMDB) and the PDB: AE1-Apo (EMD-XXXXX and XXX), AE1-Bicarbonate (EMD-XXXXX and XXX), AE1-DIDS (EMD-XXXXX and XXX), AE1-H_2_DIDS (EMD-XXXXX and XXX), AE1-DEPC (EMD-XXXXX and XXX), AE1-Dipyridamole (EMD-XXXXX and XXX), AE1-NIF (EMD-XXXXX and XXX).

## Author Information

Correspondence and requests for materials should be addressed to.

## Competing interests

The authors declare no competing interests

---

**Extended Data Fig. 1.**
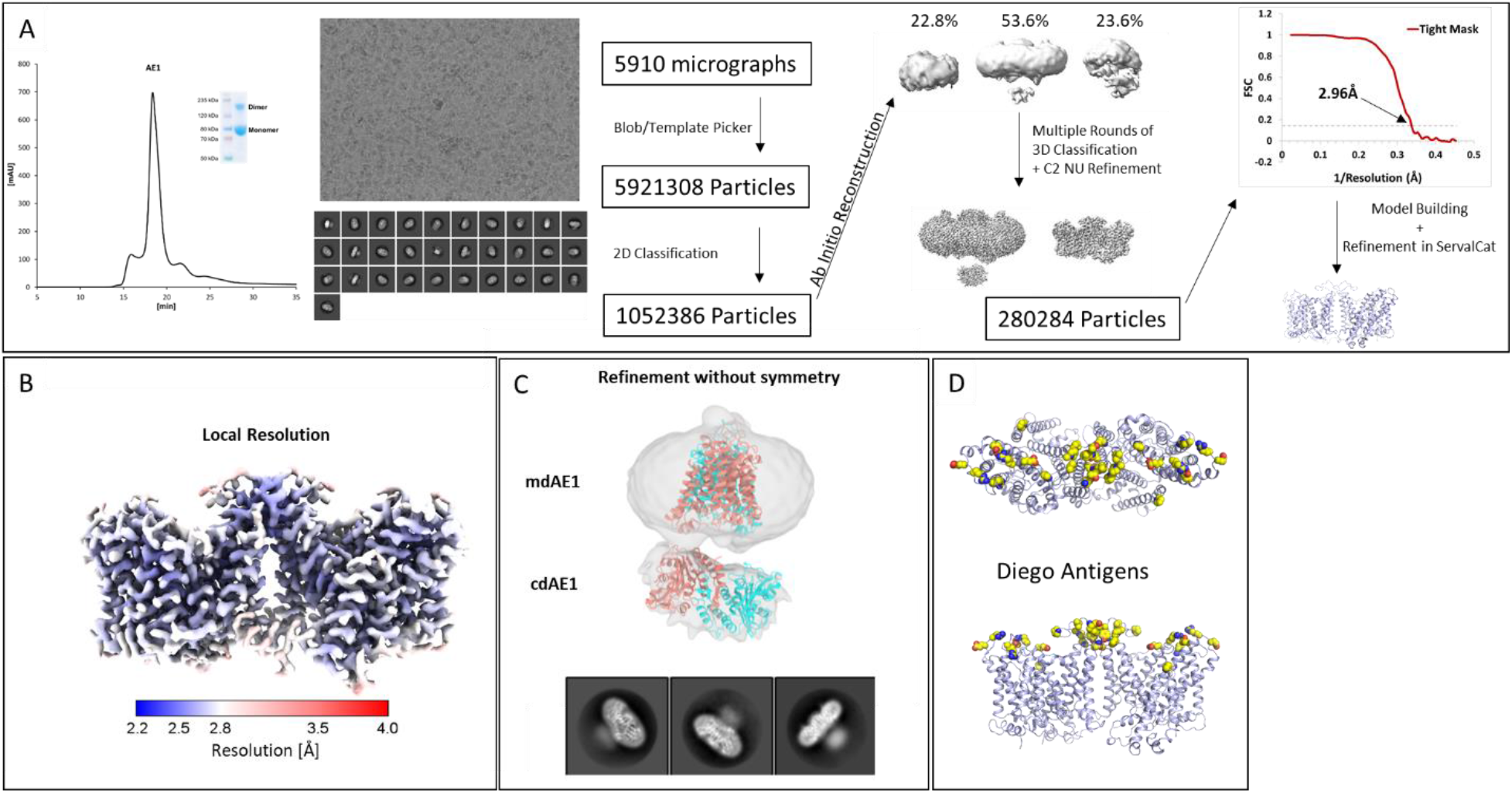
Workflow of cryo-EM Structure Determination of AE1-DIDS. **A,** Analytical size exclusion chromatography and SDS-PAGE show monodisperse and pure protein. Data were collected and processed in cryoSPARC: particles were picked from micrographs, subjected to 2D classification, followed by ab initio 3D classification. After multiple rounds of 3D classification, the final particle stack was refined using non-uniform refinement with imposed C2 symmetry. Final map was obtained with GS-FSC indicating a resolution of 2.96 Å applying the 0.143 cutoff. Initial model was built in PHENIX, and then further refined in ServalCat for the generation of final maps and coordinates of mdAE1. **B,** Calculations in cryoSPARC indicate local resolutions of up to 2.5 Å around substrate and inhibitor binding sites. **C,** Example map produced through refinement without symmetry shows tilt between cdAE1 and mdAE1, highlighting the flexibility of cdAE1. 2D classes highlight well-ordered mdAE1 density, and poorly defined cdAE1 density, respectively. **D,** Cryo-EM structures allowed us to build the complete extracellular surface including all Diego antigens (yellow).

**Extended Data Fig. 2.**
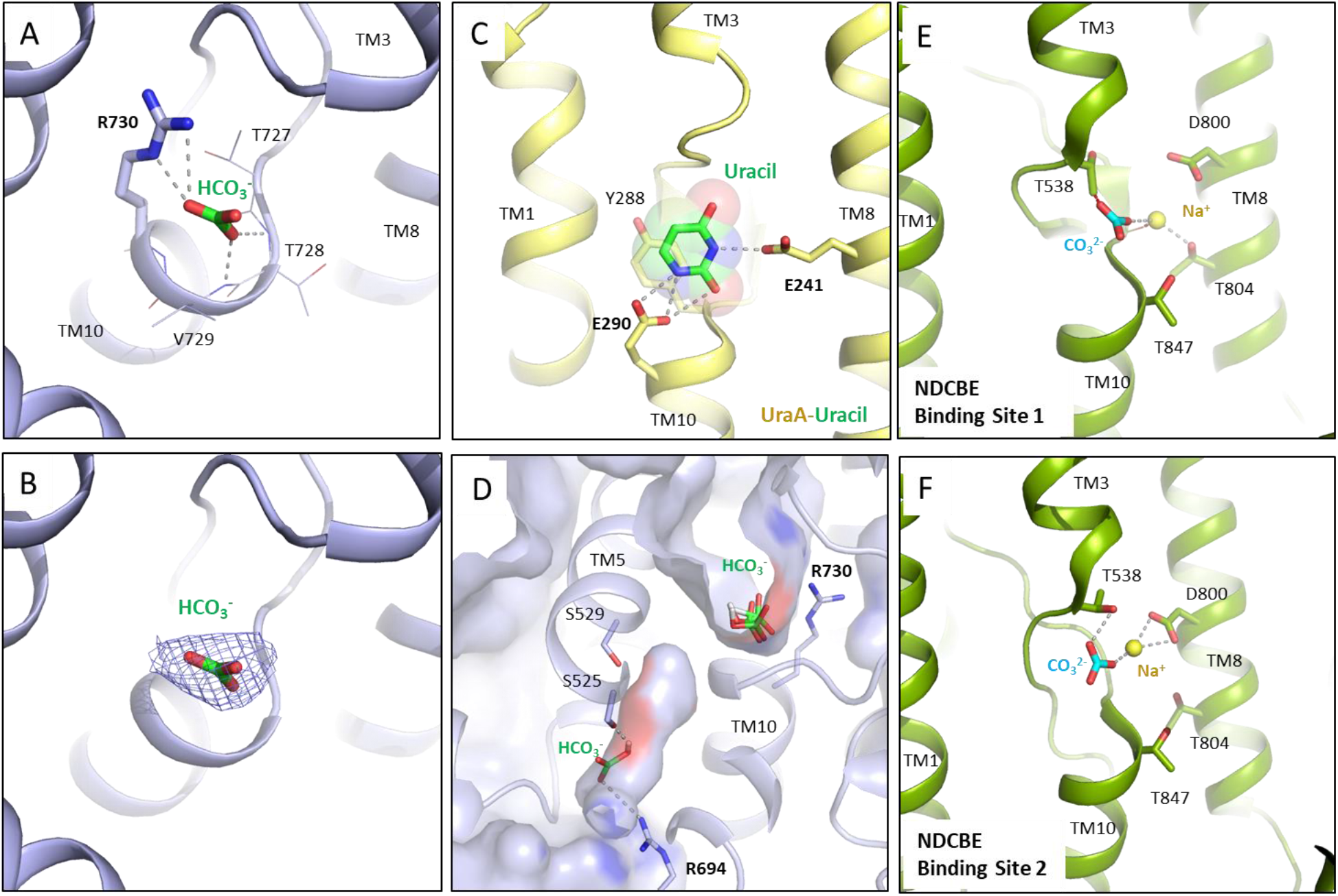
Substrate-binding in SLC4 and SLC26 transporters. **A,B,** Binding site and cryo-EM density of bicarbonate in AE1-bicarbonate structure using a contour level of 6σ. **C,** Binding site of uracil in the UraA-Uracil structure (PDB ID: 3QE7). **D,** MD simulations allow to calculate bicarbonate affinity to AE1 anion binding site and identify a second bicarbonate binding site in the putative exit channel. **E,F,** Carbonate and sodium ions in the rabbit NDCBE (SLC4A8) cryo-EM structure (PDB ID: 7RTM, EMD-24683), with clashes between CO_3_^2-^ and T538, as well as Na^+^ and the backbone shown as red dotted lines. Ionic interactions and hydrogen bonds are shown as grey dotted lines.

**Extended Data Fig. 3.**
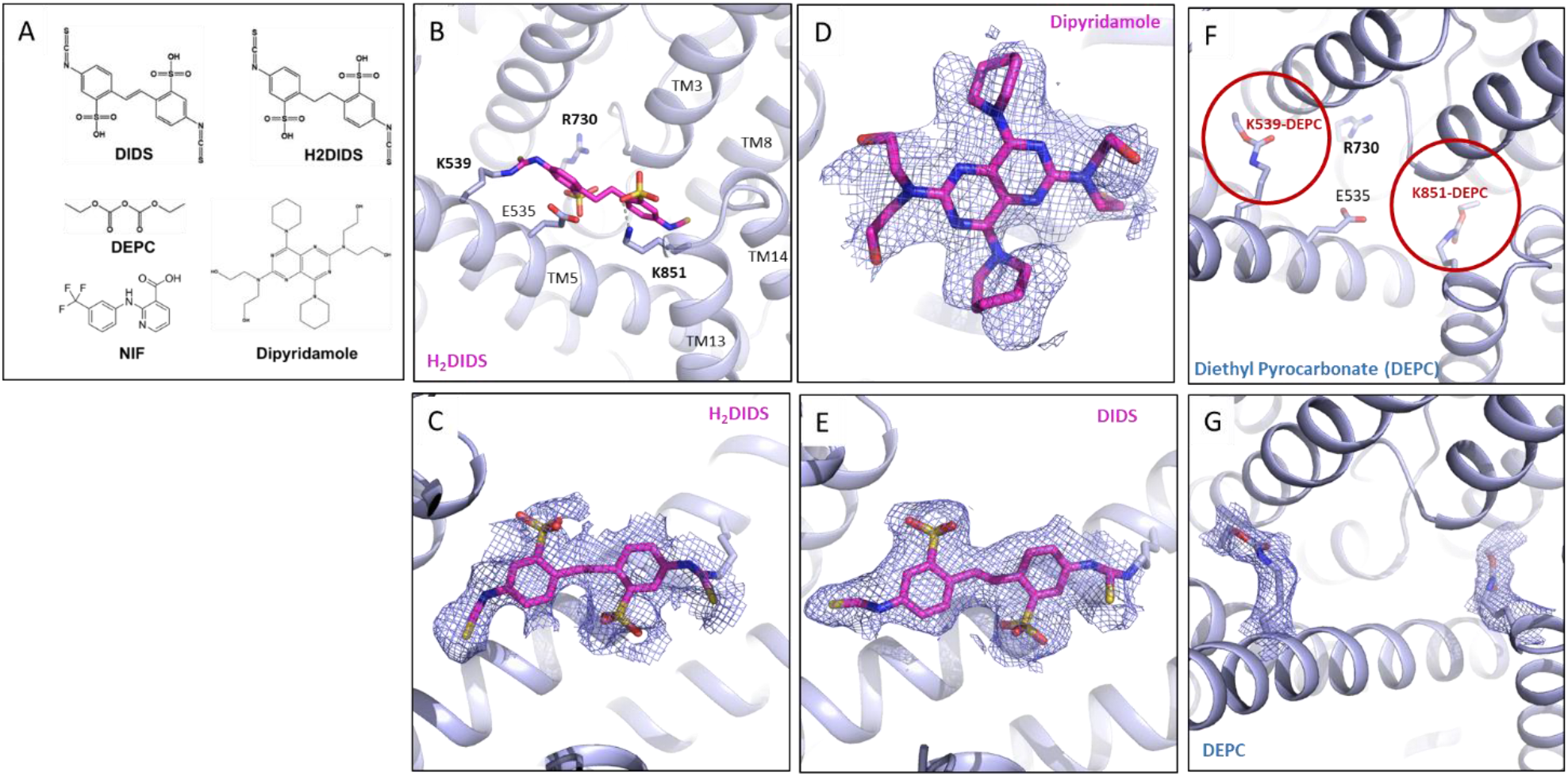
Structure and density of AE1-bound inhibitors. **A,** 2D structures of different AE1 inhibitors and ligands used in this study. **B, C,** H_2_DIDS (magenta) location and cryo-EM density at a contour level of 3σ. **D,** Cryo-EM density of Dipyridamole (contour level of 4σ). **E,** Cryo-EM density of DIDS (contour level of 3.5σ). **F, G,** DEPC modifications of K539 and K851 and cryo-EM densities shown at a contour level of 4σ.

**Extended Data Fig. 4.**
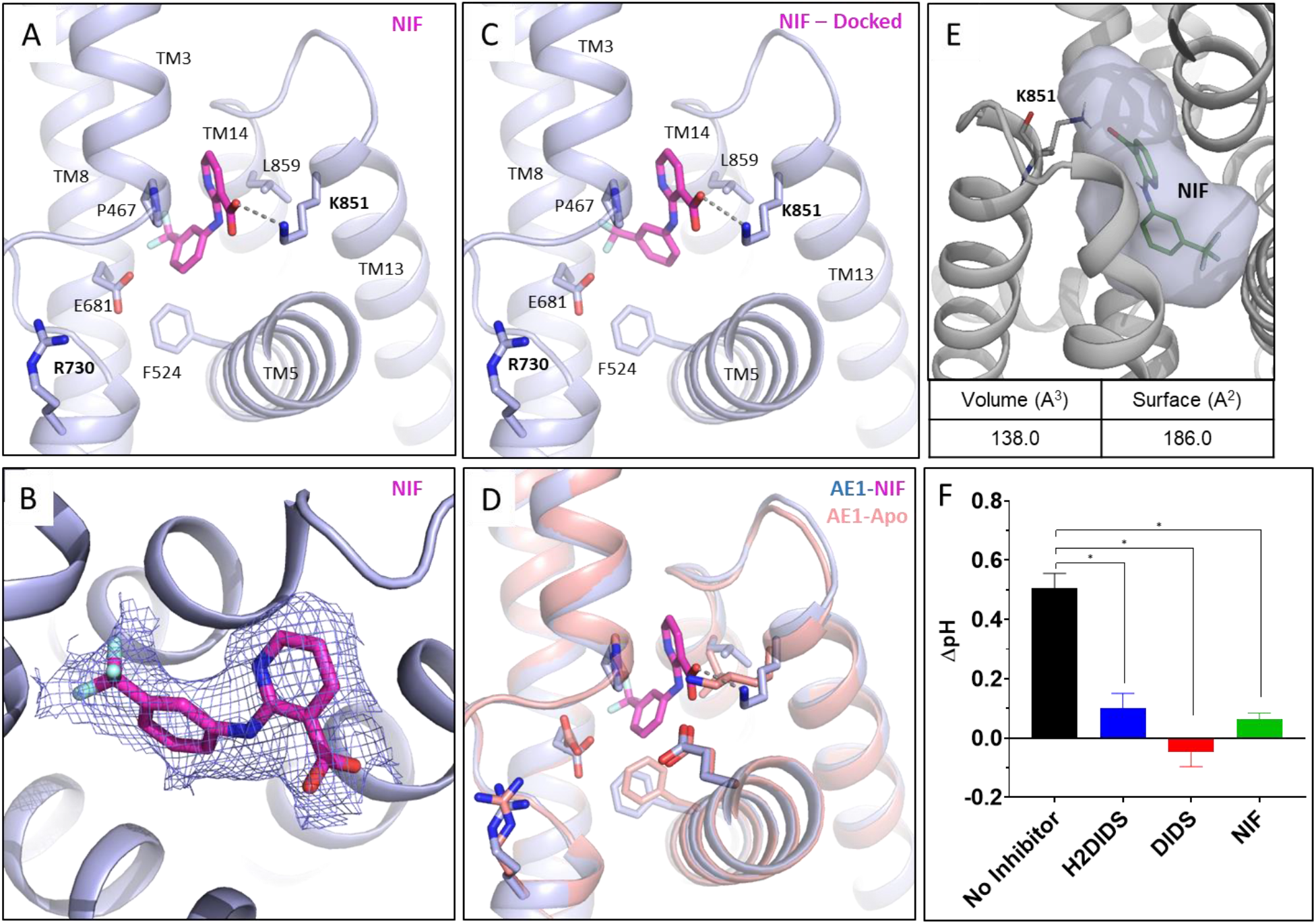
NIF binding and experimental validation of AE1-inhibition by H_2_DIDS, DIDS, and NIF. **A, B,** NIF binding site and cryo-EM density shown at a contour level of 3.5σ. **C,** Validation of NIF binding by molecular docking, with the best scoring docking pose shown (ΔG binding score: −52.33). **D**, Overlay of Apo (salmon) and NIF-bound AE1 (light blue) reveals subtle conformational rearrangements required for NIF binding. **E,** Calculation of NIF binding pocket volume and surface in POVME. Ionic interactions are shown as dotted lines. **F,** Determination of cellular bicarbonate uptake in the absence and presence of inhibitors (see methods for details). Dipyridamole could not be tested due to spectral overlap with the dye used to measure cellular pH changes. Data are mean ± s.e.m. of three independent experiments (n=3) performed in triplicates. One-way ANOVA was used to compare inhibited and uninhibited uptake (p<0.0001).

**Extended Data Table 1.**
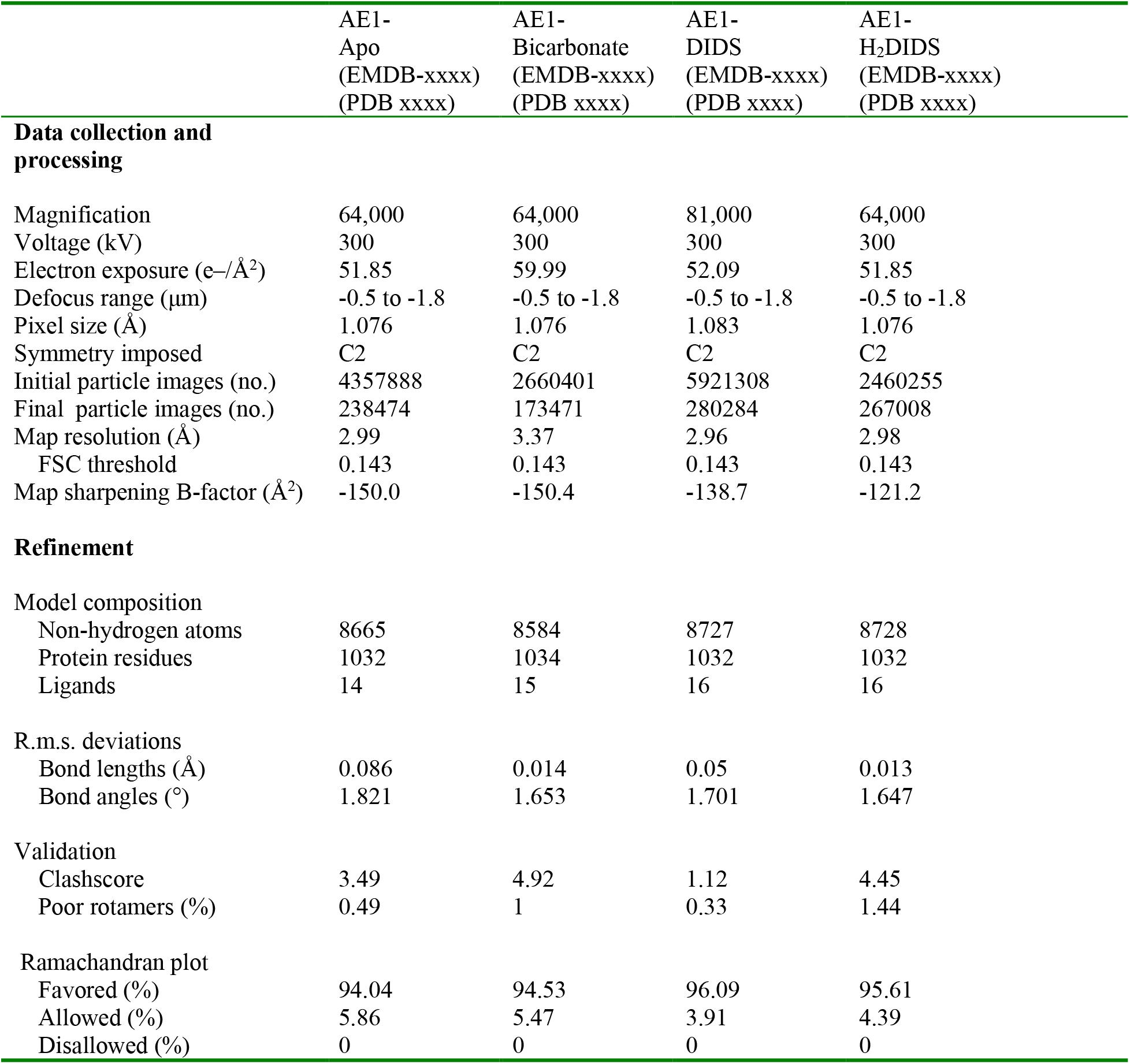
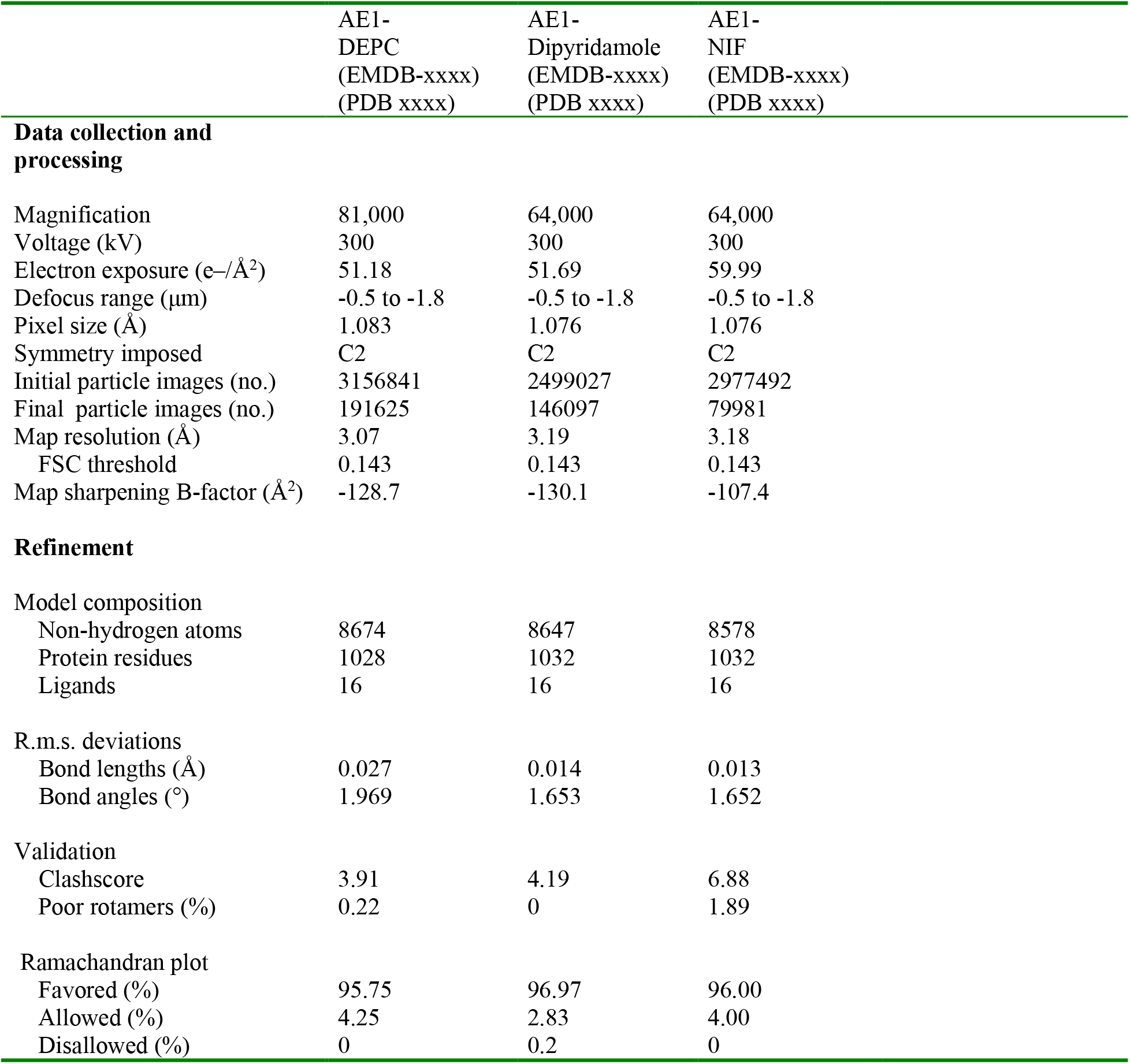
Cryo-EM data collection, refinement and validation statistics.

statistics.
|  | AE1-Apo<br>(EMDB-xxxx)<br>(PDB xxxx) | AE1-Bicarbonate<br>(EMDB-xxxx)<br>(PDB xxxx) | AE1-DIDS<br>(EMDB-xxxx)<br>(PDB xxxx) | AE1-<br>H<sub>2</sub>DIDS<br>(EMDB-xxxx)<br>(PDB xxxx) |
| --- | --- | --- | --- | --- |
| **Data collection and processing** |  |  |  |  |
| Magnification | 64,000 | 64,000 | 81,000 | 64,000 |
| Voltage (kV) | 300 | 300 | 300 | 300 |
| Electron exposure (e-/Å<sup>2</sup>) | 51.85 | 59.99 | 52.09 | 51.85 |
| Defocus range (µm) | -0.5 to -1.8 | -0.5 to -1.8 | -0.5 to -1.8 | -0.5 to -1.8 |
| Pixel size (Å) | 1.076 | 1.076 | 1.083 | 1.076 |
| Symmetry imposed | C2 | C2 | C2 | C2 |
| Initial particle images (no.) | 4357888 | 2660401 | 5921308 | 2460255 |
| Final particle images (no.) | 238474 | 173471 | 280284 | 267008 |
| Map resolution (Å) | 2.99 | 3.37 | 2.96 | 2.98 |
| FSC threshold | 0.143 | 0.143 | 0.143 | 0.143 |
| Map sharpening B-factor (Å<sup>2</sup>) | -150.0 | -150.4 | -138.7 | -121.2 |
| **Refinement** |  |  |  |  |
| Model composition |  |  |  |  |
| Non-hydrogen atoms | 8665 | 8584 | 8727 | 8728 |
| Protein residues | 1032 | 1034 | 1032 | 1032 |
| Ligands | 14 | 15 | 16 | 16 |
| R.m.s. deviations |  |  |  |  |
| Bond lengths (Å) | 0.086 | 0.014 | 0.05 | 0.013 |
| Bond angles (°) | 1.821 | 1.653 | 1.701 | 1.647 |
| Validation |  |  |  |  |
| Clashscore | 3.49 | 4.92 | 1.12 | 4.45 |
| Poor rotamers (%) | 0.49 | 1 | 0.33 | 1.44 |
| Ramachandran plot |  |  |  |  |
| Favored (%) | 94.04 | 94.53 | 96.09 | 95.61 |
| Allowed (%) | 5.86 | 5.47 | 3.91 | 4.39 |
| Disallowed (%) | 0 | 0 | 0 | 0 |

**Extended Data Table 1 (Cont) | Cryo-EM data collection, refinement and validation statistics.**
|  | AE1-DEPC<br>(EMDB-xxxx)<br>(PDB xxxx) | AE1-Dipyridamole<br>(EMDB-xxxx)<br>(PDB xxxx) | AE1-NIF<br>(EMDB-xxxx)<br>(PDB xxxx) |
| --- | --- | --- | --- |
| **Data collection and processing** |  |  |  |
| Magnification | 81,000 | 64,000 | 64,000 |
| Voltage (kV) | 300 | 300 | 300 |
| Electron exposure (e-/Å<sup>2</sup>) | 51.18 | 51.69 | 59.99 |
| Defocus range (µm) | -0.5 to -1.8 | -0.5 to -1.8 | -0.5 to -1.8 |
| Pixel size (Å) | 1.083 | 1.076 | 1.076 |
| Symmetry imposed | C2 | C2 | C2 |
| Initial particle images (no.) | 3156841 | 2499027 | 2977492 |
| Final particle images (no.) | 191625 | 146097 | 79981 |
| Map resolution (Å) | 3.07 | 3.19 | 3.18 |
| FSC threshold | 0.143 | 0.143 | 0.143 |
| Map sharpening B-factor (Å<sup>2</sup>) | -128.7 | -130.1 | -107.4 |
| **Refinement** |  |  |  |
| Model composition |  |  |  |
| Non-hydrogen atoms | 8674 | 8647 | 8578 |
| Protein residues | 1028 | 1032 | 1032 |
| Ligands | 16 | 16 | 16 |
| R.m.s. deviations |  |  |  |
| Bond lengths (Å) | 0.027 | 0.014 | 0.013 |
| Bond angles (°) | 1.969 | 1.653 | 1.652 |
| Validation |  |  |  |
| Clashscore | 3.91 | 4.19 | 6.88 |
| Poor rotamers (%) | 0.22 | 0 | 1.89 |
| Ramachandran plot |  |  |  |
| Favored (%) | 95.75 | 96.97 | 96.00 |
| Allowed (%) | 4.25 | 2.83 | 4.00 |
| Disallowed (%) | 0 | 0.2 | 0 |

**Extended Data Table 2.**
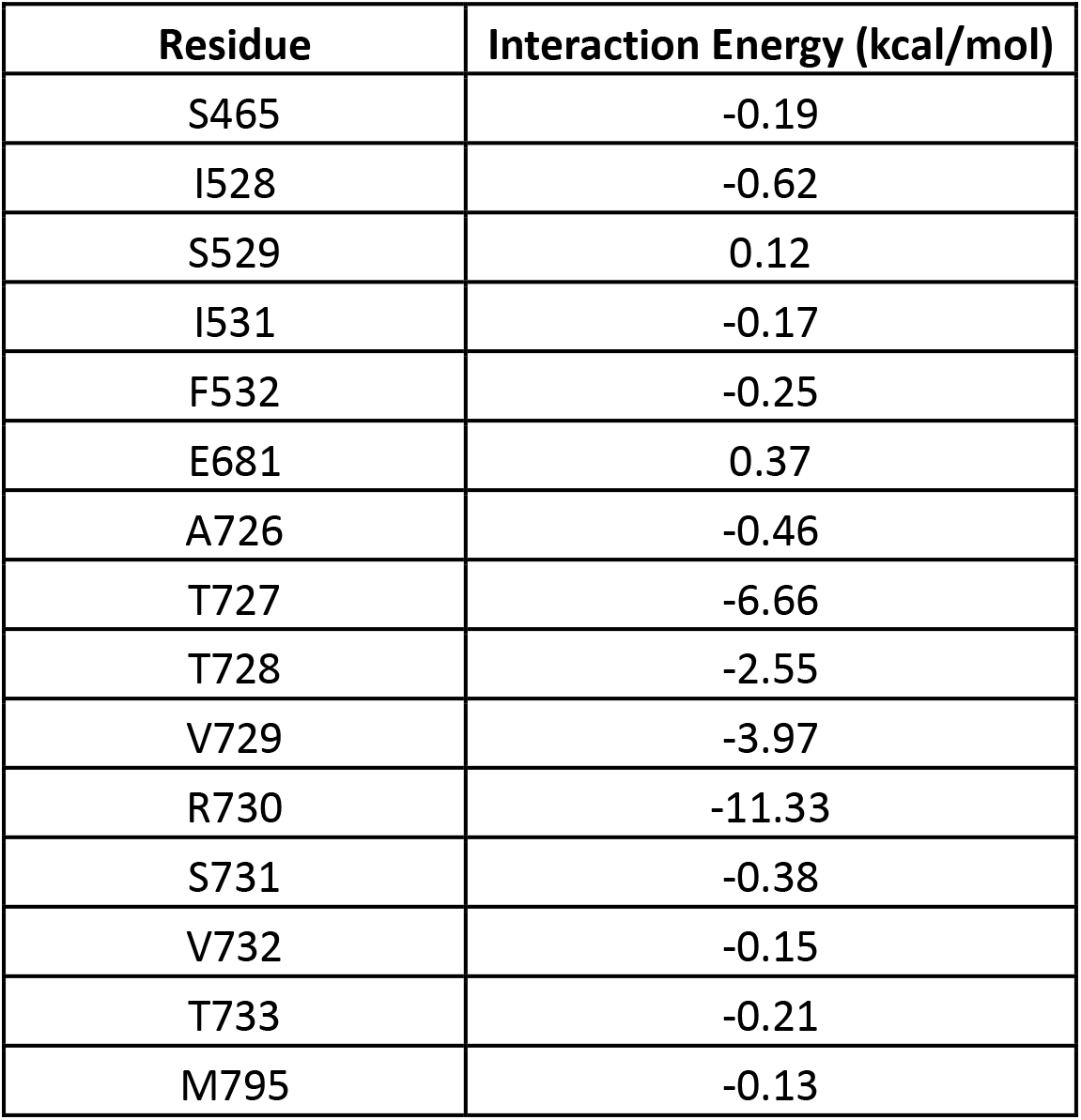
Calculated bicarbonate binding energies for residues in the AE1 anion binding site.

**Extended Data Table 3.** Calculated pocket volume of bicarbonate and inhibitors bound to AE1.

| Ligand | Volume (Å<sup>3</sup>) | Surface (Å<sup>2</sup>) |
| --- | --- | --- |
| Bicarbonate | 23 | 32 |
| H<sub>2</sub>DIDS | 312 | 341 |
| DIDS | 350 | 331 |
| Dipyridamole | 447 | 373 |
| Niflumic Acid | 138 | 186 |

## Notes

### Competing Interest Statement

The authors have declared no competing interest.

